# Binding mechanisms of therapeutic antibodies to human claudin-18.2

**DOI:** 10.64898/2026.11.09.765403

**Authors:** Xiaodong Wang, Chunting Fu, Haiyun Ma, Penghui Yang, HongYang Yu, Ziyi Sun, Liang Jin, Zhaoming Su, Xiaoming Zhou

## Abstract

Claudin-18.2 (CLDN18.2) is a tight junction component with highly restricted expression in the paracellular space of nonmalignant gastric epithelial cells. It becomes accessible on tumor cell surface during malignant transformation, making CLDN18.2 an attractive target for targeted therapies against gastric cancer. Recently, zolbetuximab has become the first FDA-approved monoclonal antibody that targets CLDN18.2 in advanced gastric cancer, and others like osemitamab have entered phase 3 clinical trials. However, the high-resolution structure of CLDN18.2 is lacking, and how CLDN18.2 is recognized by therapeutic antibodies remains undefined. Here, we report cryo-EM structures of human claudin-18.2 (hCLDN18.2) in complex with the Fab fragments of zolbetuximab and osemitamab, revealing the structure of hCLDN18.2 and its binding mechanism to the therapeutic antibodies. Further analyses reveal that three residues in the first extracellular loop of hCLDN18.2 that are different from human claudin-18.1 (hCLDN18.1) play a critical role in antibody binding. Our data elucidate the structural basis for selective binding of therapeutic antibodies to hCLDN18.2, which should aid rational design of next-generation immunotherapies targeting hCLDN18.2.

## Introduction

Gastric cancer, including gastroesophageal junction (GEJ) cancer, is one of the most prevalent malignancies worldwide(*1–4*). It is the third leading cause of cancer-related mortality globally, accounting for 7.7% of all cancer deaths(*2*). Early-stage gastric/GEJ cancer is typically asymptomatic, and the majority of patients are diagnosed at advanced stages. For these patients, primary treatments include surgery, chemotherapy, and targeted therapies(*1, 3*). For targeted therapies, human epidermal growth factor receptor 2 (HER2) has emerged as a critical therapeutic target(*1, 3, 4*). For instance, the HER2-targeting antibody-drug conjugate (ADC) trastuzumab deruxtecan (T-DXd) was approved to treat HER2-positive solid cancers, including gastric cancer(*3, 4*). However, HER2-positive gastric cancer cases constitute only 10∼20% of all gastric cancer patients, with fewer than 25% of patients benefiting from this therapy(*3, 4*). Therefore, identification of novel therapeutic targets to complement the HER2 profile for gastric cancer remains a high priority for improving clinical outcomes(*5*).

Claudins (CLDNs) are primary components of tight junctions that gatekeep the paracellular permeability of epithelial cell sheets, and contain 27 family members in mammals (26 in human)(*6–8*). Among them, the *CLDN18* gene encodes two isoforms through alternative splicing(*9*), claudin-18.1 (CLDN18.1) and claudin-18.2 (CLDN18.2), which only differ at the N-terminal 69 residues (including the first extracellular loop, ECL1)(*9–11*). CLDN18.1 is predominantly expressed in normal lung tissues, whereas CLDN18.2 is highly restricted in the paracellular space of nonmalignant gastric epithelial cells(*10*). Importantly, CLDN18.2 becomes accessible on cancer cell surface in various malignancies, including gastric, pancreatic, and esophageal cancers, making CLDN18.2 an attractive therapeutic target for treating these cancers(*10*).

The therapeutic potential of CLDN18.2 on gastric/GEJ/pancreatic cancer has spurred development of a range of anti-cancer agents, including monoclonal antibodies, ADCs, bispecific antibodies, and chimeric antigen receptor T (CAR-T) cells(*11, 12*). For instance, zolbetuximab (previously known as IMAB362 or claudiximab), a chimeric monoclonal antibody targeting human claudin-18.2 (hCLDN18.2), became the first FDA-approved treatment in combination with fluoropyrimidine-and platinum-containing chemotherapy for patients with locally advanced unresectable or metastatic HER2-negative CLDN18.2-positive gastric/GEJ adenocarcinoma in 2024(*11, 12*). A humanized antibody, osemitamab (also known as TST001), exhibits high affinity to hCLDN18.2 and has entered phase 3 trials in combination with nivolumab and chemotherapy as first-line therapy for patients with advanced or metastatic gastric/GEJ adenocarcinoma (NCT06093425). Meanwhile, more therapeutic agents targeting hCLDN18.2 are currently undergoing clinical or preclinical evaluations(*11*). However, the structural information of hCLDN18.2 is lacking, and the binding mechanisms of therapeutic antibodies to hCLDN18.2 remain undefined.

In this study, we solved the structures of hCLDN18.2 in complex with the Fab fragments of zolbetuximab and osemitamab by cryogenic electron microscopy (cryo-EM). These structures identify key residues in hCLDN18.2 ECL1 for its isoform-specific recognition by therapeutic antibodies. Combined with mutagenesis, binding analysis, and molecular dynamics (MD) simulations, our data elucidate the structural basis for selective binding of therapeutic antibodies to hCLDN18.2, providing a structural framework for rational design of next-generation immunotherapies targeting hCLDN18.2.

## Results

### Cryo-EM analysis of hCLDN18.2 bound to therapeutic antibodies

Zolbetuximab and osemitamab were selected to investigate binding of therapeutic antibodies to hCLDN18.2. Since binding occurs between the variable region (Fv, including V_H_ and V_L_) of an antibody and its antigen (Fig. S1A), we expressed the Fab fragments (containing Fv) of these two antibodies in bacterial cells (*Escherichia coli*). Subsequent purification and gel filtration analysis showed that the Fab fragments of zolbetuximab and osemitamab displayed reasonable expression and stability (Fig. S1B and S1C). Meanwhile, expression of hCLDN18.2 in yeast cells (*Pichia pastoris*) yielded high-purity hCLDN18.2 protein, which displayed two peaks in gel filtration analysis, likely representing a monomer species and a dimer species, respectively (Fig. S1D). Both peak fractions were mixed with purified Fabs for cryo-EM single-particle analysis. In the end, three complex structures, including a monomeric hCLDN18.2-zolbetuximab Fab complex (Fig. S1E), a monomeric hCLDN18.2-osemitamab Fab complex (Fig. S1F), and a dimeric hCLDN18.2-osemitamab Fab complex (Fig. S1G), were determined to 4.1 Å, 3.8 Å and 4.4 Å, respectively (Table S1).

### Molecular architecture of hCLDN18.2

Claudins can be grouped into classic or non-classic ones according to the degree of sequence similarity(*6, 13*). Classic claudins (claudin-1 to -10, -14, -15, -17, -19) share high levels of sequence homology and a common mechanism for tight junction strand assembly. In contrast, non-classic claudins (claudin-11, -12, -13, -16, -18, -20 to -27) show much higher sequence variability. Structures for several classic claudins, including mouse claudin-3, human claudin-4, human claudin-9, mouse claudin-15, and mouse claudin-19, have been reported in recent years(*14*), whereas no structure is available for a non-classic claudin (e.g. claudin-18) to date. Therefore, the hCLDN18.2-Fab complex structures in this study provide an opportunity to analyze the three-dimensional structure of a non-classic claudin (i.e. hCLDN18.2). Since the hCLDN18.2 portion of the three hCLDN18.2-Fab complex structures are largely the same with all-atom root mean square deviations (RMSDs) of < 1 Å between each other, we chose the hCLDN18.2 structure from the monomeric hCLDN18.2-osemitamab Fab complex (the highest resolution of the three) to describe its molecular architecture (Fig. S1F).

Conforming to the “claudin fold”(*15*), the hCLDN18.2 structure harbors a tetra-spanning transmembrane domain (TMD), forming a left-handed four-helix bundle (Fig. 1A). The extracellular region of hCLDN18.2 also consists of two extracellular loops (ECLs), ECL1 and ECL2, which form a five-stranded anti-parallel β-sheet (β1 to β5) as seen in the classic claudin structures(*14, 15*). ECL1 connects transmembrane helix 1 (TM1) and TM2 and contains β1 to β4, whereas ECL2 connects TM3/TM4 and contains the β5 strand. The hCLDN18.2 ECL1 also harbors a claudin consensus motif, W_30_-L_49_W_50_-C_53_-C_63_, and an intramolecular disulfide bond forms between Cys53 from β3 and Cys63 from β4 (Fig. 1A). Due to resolution limit and loop flexibility, several loops of hCLDN18.2, including the N-terminus (residues 1-3), part of ECL1 (residues 68-72) and ECL2 (residues 149-166), the intracellular loop (residues 101-113), and the C-terminus (residues 195-208), were not modeled. Aside from these unmodeled loop regions, the hCLDN18.2 structure aligns well with available claudin structures with Cα RMSDs of 1.0-1.2 Å, indicating that the overall architecture of hCLDN18.2 largely resembles the general “claudin fold” (Fig. 1B).

**Figure 1.**
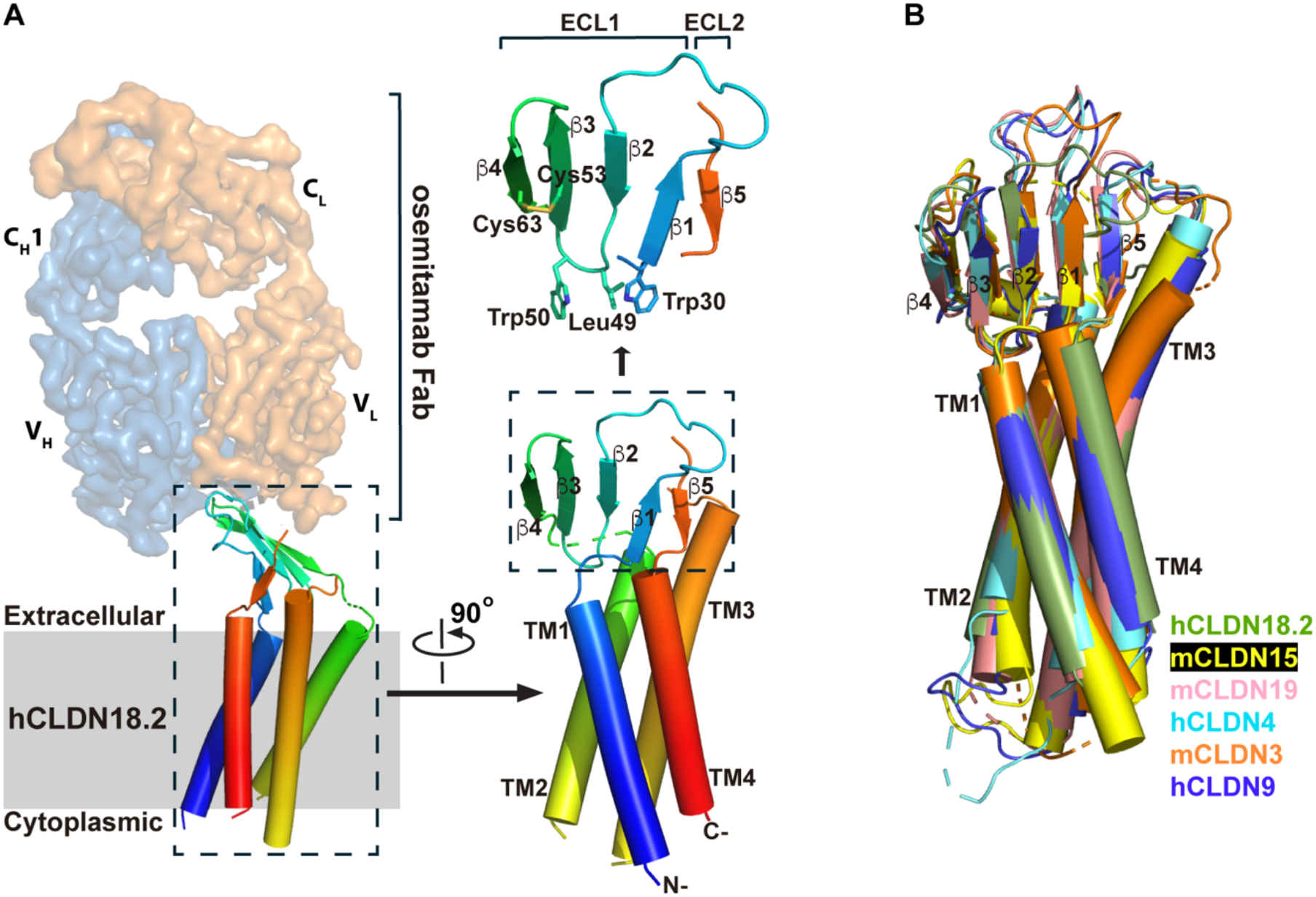
Structure of hCLDN18.2. (A) Architecture of hCLDN18.2. Left, hCLDN18.2 in the cryo-EM structure of the monomeric hCLDN18.2-osemitamab Fab complex. The osemitamab Fab is shown as cryo-EM densities in surface mode, with its light chain and heavy chain colored in orange and in blue, respectively. The hCLDN18.2 structure is displayed in cartoon mode. The membrane is indicated by a grey rectangle. Right, structural features of hCLDN18.2. The transmembrane helices (TM1-TM4) and β strands (β1-β5) are labeled. The claudin consensus motif (Trp30-Leu49-Trp50-Cys53-Cys63) and an intramolecular disulfide bond (Cys53-Cys63) are shown in sticks. (B) Superposition of hCLDN18.2 (in green) onto the structures of mCLDN15 (PDB: 4P79, in yellow), mCLDN19 (PDB: 3X29, in pink), hCLDN4 (PDB: 5B2G, in cyan), mCLDN3 (PDB: 6AKE, in orange), and hCLDN9 (PDB: 6OV2, in blue).

Besides, it is noted that the two hCLDN18.2 protomers in the dimeric hCLDN18.2-osemitamab Fab complex structure are arranged in an anti-parallel manner, with the dimer interface formed mainly between the two TMDs (Fig. S1G). Opposite topologies of the two hCLDN18.2 protomers suggest that the dimeric organization is likely a purification artefact rather than a physiological interaction, as seen in previous crystal packing of mouse claudin-19 (PDB: 3X29), human claudin-4 (PDB: 5B2G) and mouse claudin-3 (PDB: 6AKE). Therefore, the structure of dimeric hCLDN18.2-osemitamab Fab complex was not further discussed in this study.

### Binding of zolbetuximab/osemitamab Fab to hCLDN18.2

Next, the interaction between zolbetuximab/osemitamab Fab and hCLDN18.2 was analyzed. A Fab fragment contains four immunoglobulin domains, including two variable domains (V_H_ and V_L_, i.e. Fv) and two constant domains (C_H_1 and C_L_), and the Fv region mediates interactions with the antigen (Fig. S1A). As expected, the structures of hCLDN18.2-Fab complexes reveal that the Fab fragments of both zolbetuximab and osemitamab bind to the extracellular domain of hCLDN18.2 through their Fv regions (Fig. S2A). Superposition of hCLDN18.2-zolbetuximab Fab structure onto hCLDN18.2-osemitamab Fab structure yielded a Cα RMSD of 2.7 Å, suggesting a notable difference between the two complex structures (Fig. S2A). However, structural alignment between hCLDN18.2-zolbetuximab Fv region and hCLDN18.2-osemitamab Fv region, i.e. without C_H_1 and C_L_ domains, yielded a smaller Cα RMSD of 0.8 Å (Fig. S2B). The alignment results indicate that the binding complexes of zolbetuximab/osemitamab Fv regions to hCLDN18.2 are similar, and different positions of C_H_1/C_L_ domains have contributed to the larger RMSD when aligning the two hCLDN18.2-Fab structures. Therefore, we chose the hCLDN18.2-osemitamab Fab structure (3.8 Å) to analyze Fab/Fv binding to hCLDN18.2 protein.

The shape of osemitamab Fab resembles a two-claw grabber with the Fv region harboring the “claws” (Fig. 2A). Both “claws” (V_H_ and V_L_) contain three complementarity-determining regions (CDRs): CDR-H1 (residues 26-35), CDR-H2 (residues 50-58), CDR-H3 (residues 99-107) from V_H_, and CDR-L1 (residues 24-40), CDR-L2 (residues 56-62), CDR-L3 (residues 95-103) from V_L_, while the rest of V_H_/V_L_ are referred to as framework regions(*16*). CDRs are variable loops located at the tip of the Fv region, forming the antigen binding site. In the hCLDN18.2-osemitamab Fab structure, the extracellular region of hCLDN18.2 sticks into the pocket formed by the six CDRs of Fv, which interact mainly with ECL1 of hCLDN18.2 at multiple residues (Fig. 2A), including Val40 (hCLDN18.2)-Tyr100 (CDR-L3), Ala42 (hCLDN18.2)-Asn103 (CDR-H3), Asn45 (hCLDN18.2)-Asn31 (CDR-L1) (Fig. 2B), Glu56 (hCLDN18.2)-Gly102 (CDR-H3)/Asn103 (CDR-H3)/Tyr38 (CDR-L1)/Trp56 (CDR-L2) (Fig. 2C), Ser57 (hCLDN18.2)-Asn33 (CDR-H1)/Asn103 (CDR-H3)/Tyr100 (CDR-L3), and Ser58 (hCLDN18.2)-Tyr100 (CDR-H3) (Fig. 2D). Besides, Val43 (hCLDN18.2) forms hydrophobic interactions with Tyr38 (CDR-L1) and Tyr98 (CDR-L3) (Fig. 2E). The binding pattern shows that two short loop/strand sequences (residues 40-45 and 56-58) of hCLDN18.2 ECL1 mediate the main interactions with osemitamab Fab in the complex structure (Fig. 2B-2E). Meanwhile, the number of interactions suggest that CDR-H3, CDR-L1 and CDR-L3 make major contributions to hCLDN18.2 binding, whereas CDR-H1, CDR-L2 and CDR-H2 play minor roles in this process.

**Figure 2.**
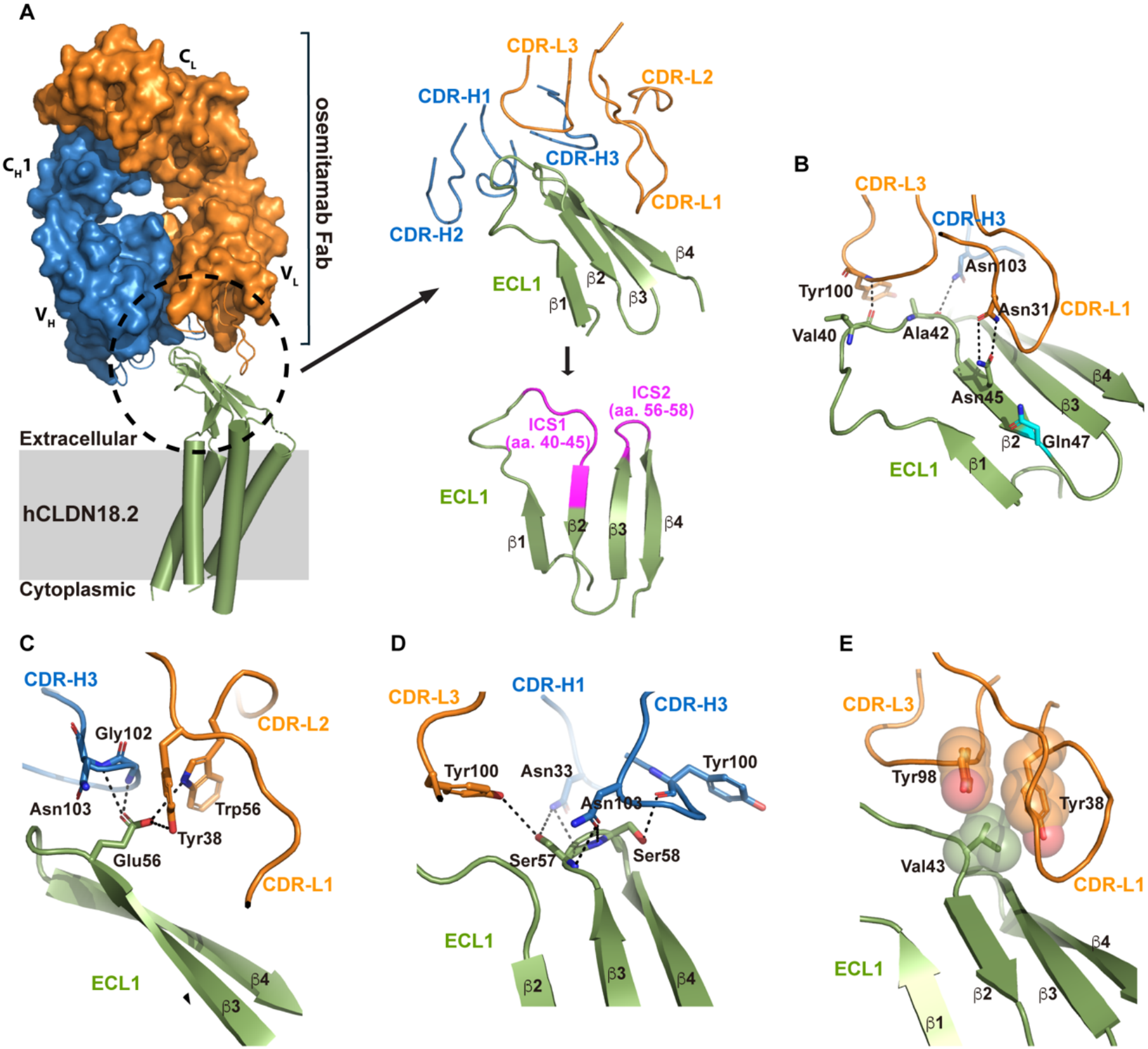
Binding of osemitamab Fab to hCLDN18.2. (A) Left, the cryo-EM structure of the hCLDN18.2-osemitamab Fab complex. Osemitamab Fab is shown in surface representation, with the heavy chain colored in blue and the light chain in orange. The hCLDN18.2 structure is displayed in green cartoon, and the membrane is indicated by a grey rectangle. The binding interface between osemitamab Fab and hCLDN18.2 is indicated by a black dashed circle. Right, close-up view of the binding interface. CDRs of the heavy and light chains of osemitamab Fab are shown as blue and orange loops, respectively. ECL1 of hCLDN18.2 is shown in green cartoon, and ICS1 and ICS2 are highlighted in magenta. (B-E) Key interactions at the binding interface between osemitamab Fab and hCLDN18.2. Participating residues are displayed as sticks (B-E) or spheres (E). Potential hydrogen bonds are indicated by black dashed lines.

In the meantime, the hCLDN18.2-zolbetuximab Fab structure displays a similar binding pattern between hCLDN18.2 ECL1 and the Fv region of zolbetuximab (Fig. S2C). Potential hydrogen bonds form between the following residue pairs: Val40 (hCLDN18.2)-Ser99 (CDR-L3), Thr41 (hCLDN18.2)-Ser99 (CDR-L3), Ala42 (hCLDN18.2)-Tyr98 (CDR-L3), Asn45 (hCLDN18.2)-Asn31 (CDR-L1) (Fig. S2D), Glu56 (hCLDN18.2)-Gly102 (CDR-H3)/Asn103 (CDR-H3)/Trp56 (CDR-L2) (Fig. S2E), Ser57 (hCLDN18.2)-Trp33 (CDR-H1)/Asn103 (CDR-H3)/Tyr100 (CDR-L3), and Ser58 (hCLDN18.2)-Asn103 (CDR-H3) (Fig. S2F). Val43 (hCLDN18.2) and Tyr38 (CDR-L1)/Tyr98 (CDR-L3) form hydrophobic interactions (Fig. S2G). In short, the main interactions also occur between residues 40-45/56-58 of hCLDN18.2 ECL1 and CDR-L1/CDR-L3/CDR-H3 of zolbetuximab (Fig. S2D-S2G), similar to the hCLDN18.2-osemitamab Fab structure with subtle differences (Fig. 2B-2E). Therefore, residues 40-45 and 56-58 of hCLDN18.2 ECL1 were designated interacting core sequence 1 (ICS1) and interacting core sequence 2 (ICS2), respectively (Fig. 2A).

### Selective binding of osemitamab Fab to hCLDN18.2 over hCLDN18.1

Although hCLDN18.2 shows a high sequence homology to hCLDN18.1 that the two differ only at the N-terminus (69 residues) (Fig. S3), hCLDN18.2-targeting therapeutic antibodies bind to hCLDN18.2, but not hCLDN18.1, specifically(*11*). The hCLDN18.2-zolbetuximab/osemitamab Fab structures show that both antibodies bind to hCLDN18.2 mainly at its ECL1 (Fig. 2A and S2C), which is located within the 69-residue N-terminus of hCLDN18.2, providing a basis for zolbetuximab/osemitamab to distinguish hCLDN18.2 from hCLDN18.1. We then attempted to solve a structure for hCLDN18.1 but did not succeed due to poor expression of hCLDN18.1. Although 21 out of the 69 N-terminal residues of hCLDN18.2 differ from that of hCLDN18.1, most differences (13 residues) occur before or within TM1 (residues 1-27) (Fig. S3), which are not accessible to therapeutic antibodies in physiological conditions and, therefore, do not interact with zolbetuximab/osemitamab. Meanwhile, residues 28-69 of hCLDN18.2 are mainly ECL1 and contains 8 differences from hCLDN18.1 (hCLDN18.1 residues in parenthesis): Gln29(Met), Asn37(Asp), Ala42(Ser), Asn45(Gln), Gln47(Glu), Glu56(Gln), Gly65(Pro), and Leu69(Ile) (Fig. 3A and S3). On the other hand, the hCLDN18.2-zolbetuximab/osemitamab Fab structures reveal that ICS1 (residues 40-45) and ICS2 (residues 56-58) of hCLDN18.2 ECL1 mediate the main interactions with zolbetuximab/osemitamab Fab (Fig. 2B-2E, and S2D-S2G), which put Ala42(Ser), Asn45(Gln) and Glu56(Gln) into focus in this study. Since Gln47(Glu) is close to ICS1 but does not interact with zolbetuximab/osemitamab Fab in the structures (Fig. 2B and S2D), we also included Gln47(Glu) in our analysis for comparison.

**Figure 3.**
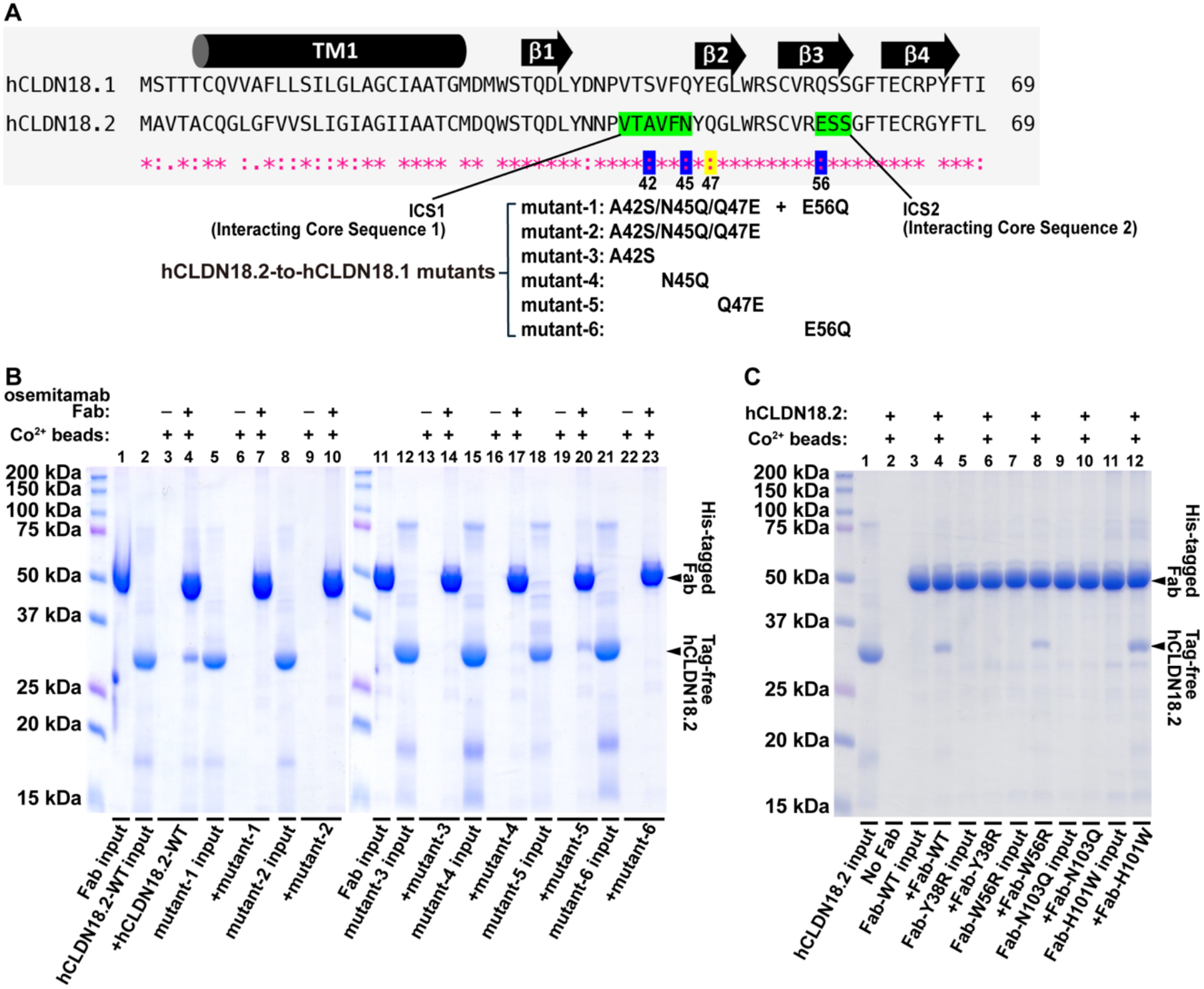
Selective binding of osemitamab Fab to hCLDN18.2 over hCLDN18.1. (A) Sequence alignment of the N-terminal 69 residues between hCLDN18.1 and hCLDN18.2. The ICS1 and ICS2 regions are highlighted in green. TM1 is indicated by a black cylinder, and β1-β4 are indicated by black arrows. Asterisks (*) indicate identical residues. Colons (:) indicate strong similarities. Periods (.) indicate weak similarities. Various hCLDN18.2-to-hCLDN18.1 mutants generated by replacing hCLDN18.2 residues with corresponding hCLDN18.1 residues are listed. (B) A pull-down assay using purified His-tagged osemitamab Fab to pull down tag-free hCLDN18.2 variants is examined by SDS-PAGE gel analysis. Signs and labels above and below the gel indicate the components present in the pull-down experiment. Molecular weight of protein standards is labeled to the left of gel. Protein bands are annotated to the right of gel. (C) A similar pull-down experiment to panel B using purified His-tagged osemitamab Fab variants to pull down tag-free wild-type hCLDN18.2.

To gain molecular insight into selective binding of therapeutic antibodies (e.g. osemitamab) to hCLDN18.2 over hCLDN18.1, we constructed a series of hCLDN18.2-to-hCLDN18.1 mutants by mutating Ala42, Asn45, Gln47 and Glu56 of hCLDN18.2 to corresponding hCLDN18.1 residues (i.e. A42S, N45Q, Q47E and Q56E) in various combinations to test their effects on antibody binding (osemitamab Fab was used here for this purpose). The designed hCLDN18.2-to-hCLDN18.1 mutants include six mutation combinations (Fig. 3A): mutant-1 (A42S/N45Q/Q47E/E56Q for ICS1+ICS2), mutant-2 (A42S/N45Q/Q47E for ICS1), mutant-3 (A42S for ICS1), mutant-4 (N45Q for ICS1), mutant-5 (Q47E near ICS1), and mutant-6 (E56Q for ICS2), which were assessed for binding to osemitamab Fab by pull-down assays. The pull-down results showed that mutant-1 (for ICS1+ICS2), mutant-2 (for ICS1) and mutant-6 (for ICS2) all lost binding to osemitamab Fab (Fig. 3B), indicating that both ICS1 and ICS2 are critical mediators for hCLDN18.2-osemitamab Fab interactions. Besides, mutant-6 contains a single mutation (E56Q), indicating that Glu56 of hCLDN18.2 is a key player in hCLDN18.2-osemitamab Fab binding. Meanwhile, single mutation analysis for ICS1 also showed that binding of mutant-3 (A42S) and, to a lesser extent, mutant-4 (N45Q) to osemitamab Fab decreased substantially. In contrast, mutant-5 (Q47E) had a mild impact on osemitamab Fab binding compared to wild-type hCLDN18.2. Consistently, mutations of Glu56-interacting or Ala42-interacting residues on osemitamab Fab (Fig. 2B and 2C, e.g. Y38R from CDR-L1, W56R from CDR-L2 and N103Q from CDR-H3 for Glu56-interacting, and N103Q from CDR-H3 for Ala42-interacting) also decreased hCLDN18.2-osemitamab Fab binding, whereas a control mutation of osemitamab Fab (H101W from CDR-H3) showed no adverse effect on its binding to hCLDN18.2 (Fig. 3C). These data demonstrate that Glu56, Ala42 and, to a lesser extent, Asn45 of hCLDN18.2 are key residues for its selective binding by osemitamab Fab over hCLDN18.1.

### Computational analysis of osemitamab binding to hCLDN18.2

Intriguingly, mutations of A42S, N45Q and E56Q of hCLDN18.2 do not seem to introduce drastic changes in binding properties of these residues in the hCLDN18.2-osemitamab Fab structure. For instance, Ala42 (hCLDN18.2) forms a hydrogen bond with Asn103 side-chain of CDR-H3 through its main-chain oxygen atom (Fig. 2B), and the A42S mutation less likely affects its main-chain atoms. Asn45 (hCLDN18.2) also forms a hydrogen bond with Asn31 of CDR-L1 via side-chain interaction (Fig. 2B), but the N45Q mutation retains hydrogen bonding ability with a slightly larger side-chain for glutamine. Meanwhile, the side-chain of Glu56 (hCLDN18.2) forms multiple hydrogen bonds with Gly102 (CDR-H3), Asn103 (CDR-H3), Tyr38 (CDR-L1) and Trp56 (CDR-L2) (Fig. 2C), whereas the E56Q side-chain is similar in size and is also capable of hydrogen bonding. These similarities in hydrogen bonding capability of residues 42, 45 and 56 between hCLDN18.2 and hCLDN18.1 makes it difficult for the hCLDN18.2-osemitamab Fab structure to self-explain why osemitamab would bind to hCLDN18.2, but not to hCLDN18.1.

Since no experimental structure for hCLDN18.2 mutants (A42S, N45Q or E56Q) in complex with antibodies (e.g. osemitamab) is available, we employed computational means to gain structural insight into mechanisms underlying the selective recognition of hCLDN18.2, but not hCLDN18.1, by osemitamab. First, Rosetta Flex ddG (*17*) was used to estimate the binding free energy change (ΔΔ*G*) between osemitamab Fab and hCLDN18.2 mutants versus wild-type hCLDN18.2. The calculated ΔΔ*G*s for mutant-1, mutant-3, mutant-4, mutant-5 and mutant-6 were 1.01, 0.28, -0.26, 0.11 and 1.33 kcal/mol, respectively (Table S2). Except for mutant-4, all other mutants caused an increase in binding free energy (Δ*G*), among which mutant-1 (A42S/N45Q/Q47E/E56Q) and mutant-6 (E56Q) showed the most prominent effects, suggesting a more severe disruption of osemitamab Fab binding to these two mutants. Therefore, the ΔΔ*G* data are largely consistent with the pull-down results.

Second, to further investigate hCLDN18.2-osemitamab Fab binding in a structural context, molecular dynamics (MD) simulations were performed for 200 ns at 298.15 K between osemitamab Fab and hCLDN18.2 variants (including wild-type, mutant-1, mutant-6, and a negative control N146Q far away from the interacting zone). Based on RMSD analysis, the last 20 ns of each trajectory were used for Δ*G* calculations using the MM/PBSA approach(*18*) (Fig. S4). The calculated Δ*G* were -73.98, -66.12, and -53.69 kcal/mol for wild-type, mutant-1, and mutant-6, respectively (Table S2), indicating significantly reduced binding affinities of osemitamab Fab for hCLDN18.2 mutant-1/mutant-6. During the 200-ns simulations, residues 56-62 (near ICS2) of hCLDN18.2 showed increased root mean square fluctuations (RMSFs) in the mutant-1/mutant-6-osemitamab Fab complexes, whereas residues 35-42 (near ICS1) exhibited increased flexibility in the mutant-6-osemitamab Fab complex (Fig. S5A). Notably, increased RMSFs were also observed in the CDR regions of osemitamab Fab, including V_H_ residues 100-104 (CDR-H3), and V_L_ residues 27-37 (CDR-L1) and 97-102 (CDR-L3) (Fig. S5B and S5C). These regions are consistent with the interacting residues identified in the hCLDN18.2-zolbetuximab/osemitamab Fab structures, suggesting that the hCLDN18.2-to-hCLDN18.1 mutations (e.g. E56Q in mutant-6) weakened the interactions between hCLDN18.2 mutant-1/mutant-6 and osemitamab Fab. In contrast, the RMSFs of the negative control (the hCLDN18.2_N146Q_-osemitamab Fab complex) remained largely unchanged in these regions. In addition, the average number of hydrogen bonds during the simulation was estimated to be 13.1 for wild-type hCLDN18.2, 11.5 for mutant-1, 10.4 for mutant-6, and 12.7 for the negative control N146Q, which also shows fewer/weaker interactions between osemitamab Fab and hCLDN18.2 mutant-1/mutant-6. Furthermore, RMSDs of the interacting residues from osemitamab Fab were calculated following least-squares fitting to the interacting residues of hCLDN18.2, which increased significantly in both mutant-1 and mutant-6 complexes compared to wild-type and the negative control complexes (Fig. S6 and Table S3). These simulation data demonstrate coherently that the hCLDN18.2-to-hCLDN18.1 mutations (e.g. A42S, N45Q and E56Q) cause disruption in the interactions between hCLDN18.2 and osemitamab Fab, providing a structural basis for selective targeting of hCLDN18.2 over hCLDN18.1 by therapeutic antibodies like osemitamab.

## Discussion

In this study, we resolved two complex structures between hCLDN18.2 and zolbetuximab/osemitamab Fab, from which key sequences/residues of hCLDN18.2 were identified that mediate interactions with the antibodies. Meanwhile, it is also desirable to analyze how antibody CDR sequences (e.g. zolbetuximab/osemitamab) affect their binding to hCLDN18.2, which may provide clues for rational design of next-generation immunotherapies targeting hCLDN18.2.

The hCLDN18.2-zolbetuximab/osemitamab Fab structures show that both zolbetuximab and osemitamab bind to hCLDN18.2 ECL1 with similar CDR regions. We then compared Fv sequences of eight hCLDN18.2-targeting antibodies (or conjugates) for clues of general CDR positions that are critical in hCLDN18.2 binding. The eight antibodies include monoclonal antibodies zolbetuximab, osemitamab and omectatug (also known as ASKB589), ADCs tecotabart vedotin, ciletatug vedotin (also known as RC118) and sonesitatug vedotin (also known as CMG901), a bispecific antibody spevatamig (also known as PT886), and a CAR-T cell antibody satricabtagene autoleucel (also known as CT041). Sequence alignment of the antibodies showed that CDR-L1, CDR-L2 and CDR-L3 are apparently less variable than CDR-H1, CDR-H2 and CDR-H3 (Fig. S7). Interestingly, the CDR residues interacting with hCLDN18.2 residues Ala42, Asn45 and Glu56 in the hCLDN18.2-zolbetuximab/osemitamab Fab structures, including Tyr98 (in CDR-L3), Asn103 (in CDR-H3), Asn31 (in CDR-L1), Gly102 (in CDR-H3), Tyr38 (in CDR-L1) and Trp56 (in CDR-L2), are invariable in all eight antibodies (Fig. S7). Potentially, these CDR positions or residues in the vicinity may serve as a start point for manipulation to tune the hCLDN18.2-antibody interactions.

## Methods

### Expression and purification of hCLDN18.2

The gene encoding the full-length hCLDN18.2 (NCBI accession NP_001002026.1) was codon-optimized for *Pichia pastoris* and synthesized (Genewiz, China). An hCLDN18.2 fragment (encoding residues 1-208) was then subcloned into a modified pPICZ plasmid (Thermo Fisher Scientific) containing an N-terminal tag of FLAG-His_10_-TEV protease recognition site for structural analysis in this study. For pull-down assays, a bacterial cytochrome b562RIL (BRIL) module was fused to the C-terminus of the hCLDN18.2 fragment in the aforementioned construct to separate protein bands on electrophoresis gels. All hCLDN18.2 mutations were introduced by QuikChange II system (Agilent) according to manufacturer’s recommendation, and were verified by DNA sequencing. The constructs were linearized and transformed into yeast strain GS115 (*Pichia pastoris*) by lithium chloride/single-strand carrier DNA/polyethylene glycol method according to manufacturer’s manual (Thermo Fisher Scientific). The transformants were inoculated into YPD medium consisting of 1% (w/v) yeast extract, 2% (w/v) peptone and 2% (w/v) D-(+)-glucose at 30 °C with shaking at 220 rpm until an optical density at 600 nm (OD_600_) of 3-5 was reached. To induce protein expression, yeast cells were harvested by centrifugation and resuspended to an OD_600_ of 1 in YPM medium consisting of 1% (w/v) yeast extract, 2% (w/v) peptone, 0.8% (v/v) methanol and 2.5% (v/v) dimethyl sulfoxide (DMSO) at 30 °C for 48 h. Cell pellets were resuspended in Lysis Solution (LS) containing 20 mM Tris-HCl pH 7.5, 150 mM NaCl, 10% (v/v) glycerol, 1 mM phenylmethanesulfonyl fluoride (PMSF), and were lysed by an AH-1500 high-pressure homogenizer (ATS, China) at 1,300 MPa. Undisrupted cells and cell debris were separated by centrifugation at 3,000 x g, and membranes were collected by ultracentrifugation at 140,000 x g for 1 h at 4 °C. The membrane pellet was resuspended into LS using a glass Dounce homogenizer and was solubilized by 1% (w/v) n-dodecyl-β-D-maltopyranoside (DDM, Anatrace) at 4 °C for 2 h and the extraction mixture was centrifuged at 200,000 x g for 30 min at 4 °C. The supernatant was then incubated with Co^2+^ resin in the presence of 20 mM imidazole pH 8.0 at 4 °C for 1 h, and the mixture was loaded in an empty chromatography column. The resin/protein was washed with 20 bed-volume of LS containing 2 mM DDM and 30 mM imidazole pH 8.0, and the protein was eluted with LS supplemented with 2 mM DDM and 250 mM imidazole pH 8.0. Affinity-purified hCLDN18.2 protein was incubated with TEV protease at room temperature for 2 h to cleave the His-tag and loaded onto a Superdex 200 Increase 10/300 GL column (GE Healthcare) for purification by size-exclusion chromatography (SEC).

### Expression and purification of Fab fragments of therapeutic antibodies

All antibody sequences used in this study were obtained from the GSRS website (FDA Global Substance Registration System, https://gsrs.ncats.nih.gov/ginas/app/ui/home). The genes encoding Fab fragments of zolbetuximab and osemitamab were codon-optimized for *Escherichia coli*, and were synthesized (Genewiz, China) and cloned into a modified pET plasmid containing two T7 promoters, which drive expression of the Fab heavy chain and light chain, respectively. The Fab heavy chain contains an N-terminal PelB signal peptide and a C-terminal TEV-FLAG-His10 tag, while the light chain contains an N-terminal PelB signal peptide. The Fab constructs were then transformed into BL21-Gold (DE3) cells. The transformants were grown in LB media at 37 °C with 50 μg/ml Kanamycin to OD_600_ of 1, followed by induction with 1 mM isopropyl β-D-1-thiogalactopyranoside (IPTG) and overnight shaking (180 rpm) at 25 °C. Cell pellets were resuspended in TES buffer (0.2 M Tris-HCl pH 8.0, 0.5 mM EDTA, 0.5 M sucrose) and were shaken at 4 °C for 2 h. Cells underwent osmotic shock by addition of an equal volume of ice-cold water and were shaken for 45 min at 4 °C. The cell pellet was discarded after centrifugation at 50,000 x g for 20 min at 4 °C. The supernatant was then loaded onto a cobalt metal affinity column, washed with wash buffer (20 mM Tris-HCl pH 7.5, 150 mM NaCl, 10% glycerol) supplemented with 5 mM imidazole pH 8.0. The Fab fragments were then eluted with wash buffer supplemented with 250 mM imidazole pH 8.0.

### Cryo-EM sample preparation and data acquisition

Recombinant hCLDN18.2 and Fab fragments were further purified by SEC in a buffer consisting of 150 mM NaCl, 20 mM Tris-Cl pH 7.5, and 0.5 mM DDM. The peaks corresponding to monomer or dimer hCLDN18.2 were collected separately, which were incubated with molar-excessive Fab fragments on ice for 30 min to form complexes. The complex samples were purified by SEC in 20 mM Tris pH 7.5, 150 mM NaCl, and 0.5 mM DDM. The peak fractions corresponding to a monomeric hCLDN18.2-zolbetuximab Fab complex, a monomeric hCLDN18.2-osemitamab Fab complex, and a dimeric hCLDN18.2-osemitamab Fab complex were collected and concentrated to 8.0 mg/ml, 11.0 mg/ml, and 8.6 mg/ml, respectively.

Purified complex samples were prepared for cryo-EM analysis. For each complex, 3.0 μl solution was applied to glow-discharged (45 s, EasGlow, PELCO) holey carbon grids (Quantifoil Au R2/1, 200 mesh). The grids were blotted for 3.0 s and plunge-frozen into liquid ethane, pre-cooled with liquid nitrogen, using a Vitrobot Mark IV system (Thermo Fisher Scientific) maintained at 4 °C and 100% humidity. Cryo-EM data were acquired on a Titan Krios cryogenic electron microscope (Thermo Fisher Scientific) operating at 300 kV. The microscope was equipped with a Gatan K2 Summit direct electron detector and a BioQuantum energy filter (Gatan) operated with a 20 eV slit. A 50 μm condenser lens aperture was used, and images were recorded at a nominal magnification of 165,000×, corresponding to a calibrated pixel size of 0.85 Å. Movie stacks were automatically collected using EPU software (Thermo Fisher Scientific) with the K2 detector operating in counting mode. The electron exposure was fractionated into 30 frames over a 6 s exposure time (5 raw frames per second), resulting in a total electron dose of approximately 58 e⁻/Å². Datasets comprising 5,883, 1,588, and 3,994 movie stacks were collected for the osemitamab-hCLDN18.2 monomer complex, osemitamab-hCLDN18.2 dimer complex, and zolbetuximab-hCLDN18.2 monomer complex, respectively. The defocus range during data collection was set from -1.0 to -1.8/-3.0 μm.

### Cryo-EM data processing and 3D reconstruction

For the osemitamab-hCLDN18.2 monomer complex, cryo-EM data processing was performed using Relion(*19*) (v5.0) and cryoSPARC(*20*) (v4.4). In total, 5,883 movies were collected. The raw movie stacks were motion corrected using Patch Motion Correction, followed by contrast transfer function (CTF) estimation using Patch CTF Estimation within cryoSPARC. Images with a CTF resolution better than 4 Å were selected, resulting in 1,729 images. A subset of the data (500 images) was used to auto-pick particles by the Blob Picker function in cryoSPARC. After 2D classification, 32,681 particles with different views were selected for template-based particle picking. In total, 843,125 particles were picked and extracted with a box size of 288 pixels and cropped to 128 pixels. After three rounds of 2D classification, ab-initio reconstruction and heterogenous refinement, a map with apparent transmembrane helix densities was generated, whose particles were re-extracted with a pixel size of 0.85 Å. To further improve the resolution, seed-facilitated 3D classification was performed. In total, 73,763 particles were imported into Relion for Blush 3D auto-refinement, followed by Bayesian polishing. Polished particles were imported back into cryoSPARC for non-uniform refinement, CTF refinement and local refinement, yielding a final density map of 3.8 Å. The resolution estimations were based on the Fourier shell correlation of 0.143 cut-off after correction of the masking effect. The B-factor used for map sharpening was automatically determined by non-uniform refinement in cryoSPARC. Local resolution maps were calculated using cryoSPARC (Fig. S8).

For the zolbetuximab-hCLDN18.2 monomer complex, 3,994 movie stacks were collected, and were processed similarly for motion correction and CTF estimation. After manual inspection, 3,925 micrographs were selected. A total of 2,725,391 particles were extracted using a box size of 256 pixels. Following two rounds of reference-free 2D classification, 721,639 particles from well-defined classes were selected for ab-initio reconstruction (5 classes) and heterogeneous refinement. The optimal 3D class, containing 300,203 particles (41.6% of the input), was subjected to homogeneous refinement, non-uniform refinement, reference-based motion correction, and local refinement, resulting in a 4.1 Å density map (Fig. S9).

For the osemitamab-hCLDN18.2 dimer complex, the 1,588 movie stacks underwent similar motion correction and CTF estimation. Manual inspection yielded 1,543 micrographs for particle picking. A total of 1,379,077 particles were extracted with a box size of 320 pixels. After two rounds of reference-free 2D classification, 386,714 particles from well-defined classes were selected for ab-initio reconstruction (6 classes) and heterogeneous refinement. The best 3D class, containing 141,022 particles (36.5% of the 2D-selected particles), was then subjected to 3D classification (3 classes). The optimal class from this step, containing 66,529 particles (47.2% of the input), was selected for homogeneous refinement, followed by non-uniform refinement, reference-based motion correction, and local refinement, yielding a final density map of 4.4 Å resolution (Fig. S10). The resolutions for all final maps were determined using the 0.143 cut-off of FSC in cryoSPARC. Final density maps were low-pass filtered according to their global resolution and visualized in UCSF Chimera(*21*).

### Cryo-EM model building and structure refinement

Cryo-EM model building was facilitated by an AlphaFold(*22*)-predicted hCLDN18.2 model (AlphaFold DB: AF-P56856-F1) and a published Fab model (PDB: 1FOR). Both hCLDN18.2 and Fab models were manually adjusted and rebuilt in Coot(*23*) and refined using the phenix.real_space_refine module(*24*) in PHENIX(*25*). Some flexible loop regions were removed due to unresolved densities. Model geometries were assessed by Molprobity(*26*) and summarized in Table S1. All structural figures, RMSD calculations, and length measurements were performed in PyMOL (Schrӧdinger, LLC). Some cryo-EM densities were visualized in UCSF Chimera(*21*).

### Sequence alignment

All sequence alignments were performed by ClustalW(*27, 28*).

### Pull down assay

For pull-down assays, affinity-purified hCLDN18.2-BRIL and variants were treated with TEV protease to remove the His-tag, and were further purified by SEC. Then, purified Fab fragments (containing the His-tag) were incubated with Co^2+^ Beads for 30 min at 4 °C, followed by mixing with His-tag-free hCLDN18.2-BRIL and variants in a final volume of 1 ml for 40 min at 4 °C. After incubation, the mixture was centrifuged at 5,000 ×g for 1 min at 4 °C to discard the supernatant. The resulting beads were washed twice with the mixing buffer before SDS-PAGE analysis.

### Molecular dynamics (MD) simulation

The mutants of the monomeric hCLDN18.2-osemitamab Fab complex were modeled, and their binding affinities were estimated using the Rosetta Flex ddG protocol(*17*). These models were subsequently subjected to molecular dynamics (MD) simulations. The non-covalent interactions were analyzed and presented by PyMOL (Schrӧdinger, LLC). MD simulation was performed by GROMACS-2024(*29*) with the Amber14sb force field(*30*). The modeling system was solvated in a cubic box filled with a TIP3P water model(*31*), maintaining a minimum distance of 15 Å between the complex and the box boundary. Then, the charges of the system were neutralized using Na^+^ or Cl^-^, followed by the steepest descent method for energy minimization. Finally, the MD simulation was conducted at 298.15 K for 200 ns after performing NVT (isochoric-isothermal) and NPT (isothermal-isovolumetric) ensembles under the same temperatures for 200 ps. The atomic trajectories were analyzed to compute the root-mean-square deviation (RMSD), root-mean-square fluctuation (RMSF) and hydrogen bonds.

## Data Availability

The cryo-EM density maps generated in this study have been deposited in the Electron Microscopy Data Bank (EMDB) under the following accession codes: EMDB-64747 (monomeric hCLDN18.2-zolbetuximab Fab complex), EMDB-64740 (monomeric hCLDN18.2-osemitamab Fab complex), and EMDB-64746 (dimeric hCLDN18.2-osemitamab Fab complex). Their corresponding atomic coordinates have been deposited in the Protein Data Bank (PDB) under the accession codes 9V32 (monomeric hCLDN18.2-zolbetuximab Fab complex), 9V2U (monomeric hCLDN18.2-osemitamab Fab complex), and 9V31 (dimeric hCLDN18.2-osemitamab Fab complex).

## Acknowledgements

The initial cryo-EM sample screening was conducted in Bioland Laboratory. The final cryo-EM data were collected at SKLB West China Cryo-EM Center, and were processed at the Duyu High Performance Computing Center of Sichuan University and on our in-house workstations. This work was supported in part by Sichuan Science and Technology Program grant 2023ZYD0125 to X.Z., West China Hospital of Sichuan University “1.3.5 Project” for Disciplines of Excellence grants ZYYC20014 to X.Z. and ZYYC21006 to Zh.S., and National Key Research and Development Program of China grant 2024YFF0727300 and Major Project of Guangzhou National Laboratory GZNL2023A03008 to L.J.

## Author Contributions

X.Z. and Zi.S. conceived the project and wrote the manuscript with input from all authors. X.Z., Zh.S. and L.J. designed the experiments. X.W. prepared the samples and performed functional studies. C.F., H.M. and X.W. processed cryo-EM data. P.Y. performed computational studies. Zh.S., H.M., L.J. and H.Y. performed cryo-EM sample screening and data collection. X.Z., X.W., C.F. and H.M. built structural models. All authors analyzed the data.

## Supplementary Figures and Tables

**Figure S1.**
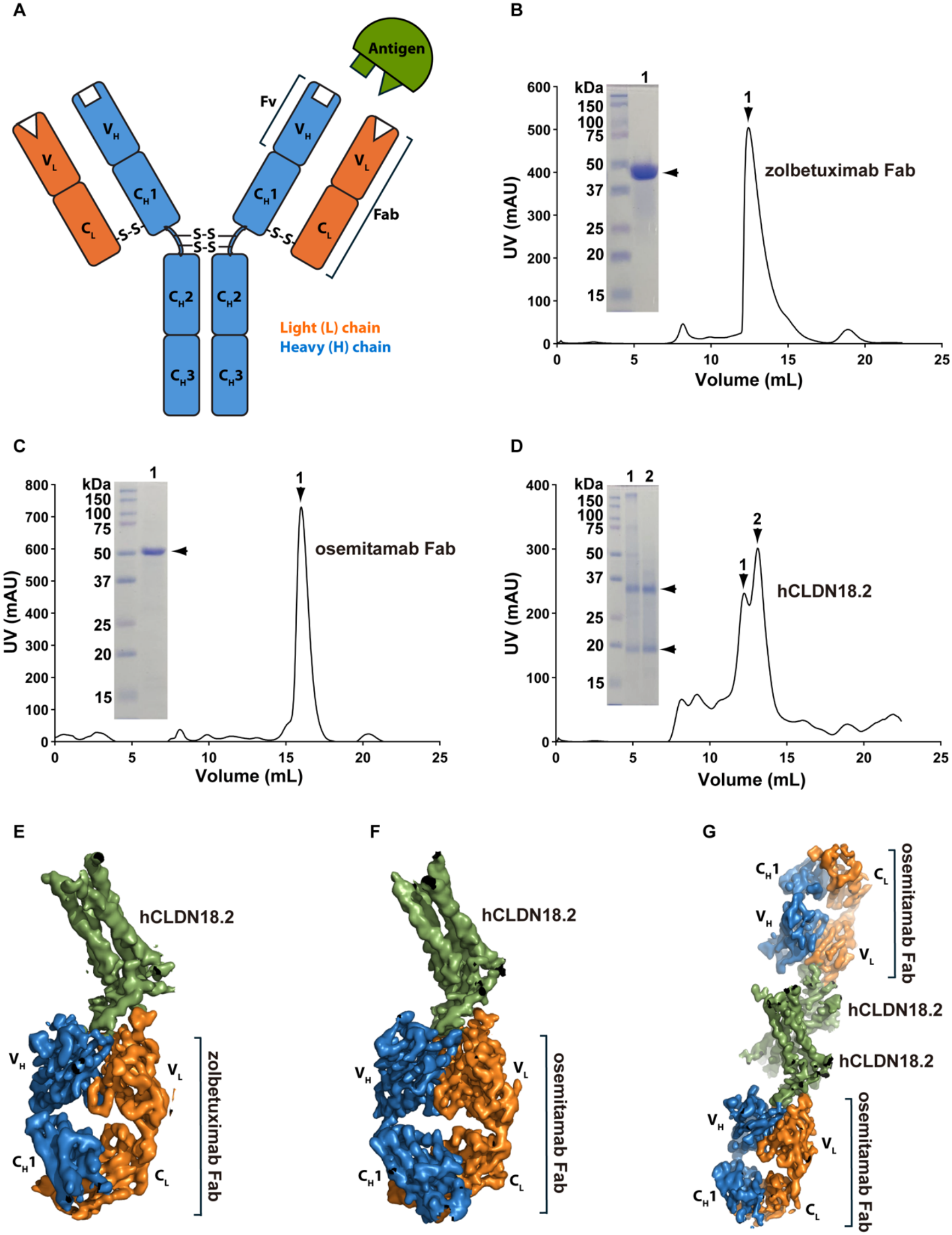
Sample preparation of complexes between hCLDN18.2 and therapeutic antibody Fab fragments. (A) Schematic diagram of an antibody and its binding to an antigen. The antibody heavy chain is shown in blue, the light chain in orange, and the antigen in green. The Fab and Fv regions are indicated. (B-C) Size-exclusion chromatography profiles and SDS-PAGE analyses of purified zolbetuximab Fab and osemitamab Fab, respectively. The arrowheads mark the peak and band positions corresponding to the Fab fragments. (D) Size-exclusion chromatography profile of purified hCLDN18.2, with peaks 1 and 2 corresponding to the dimer and monomer species, respectively. Lanes 1 and 2 of the gel indicate protein samples from peaks 1 and 2 are loaded. The monomer and dimer band positions are indicated by two arrowheads to the right of the gel. (E-G) Cryo-EM densities of a monomeric hCLDN18.2-zolbetuximab Fab complex (E), a monomeric hCLDN18.2-osemitamab Fab complex (F), and a dimeric hCLDN18.2-osemitamab Fab complex (G). The heavy and light chains of Fab fragments are shown in blue and orange, respectively, and hCLDN18.2 in green.

**Figure S2.**
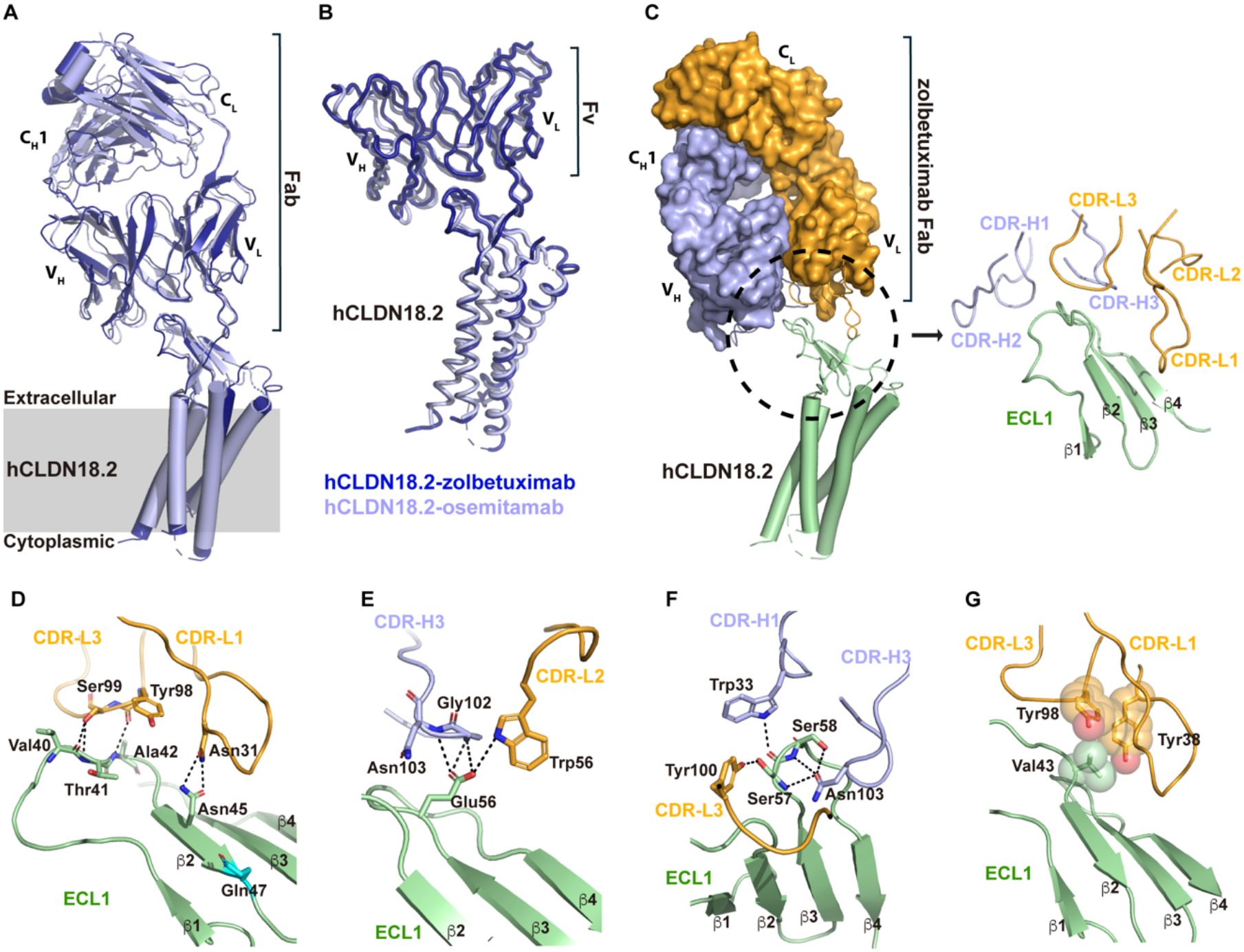
Binding of zolbetuximab Fab to hCLDN18.2. (A) Superposition of hCLDN18.2-zolbetuximab Fab structure (in dark blue) onto the structure of hCLDN18.2-osemitamab Fab (in light blue). The membrane is indicated by a grey rectangle. (B) Superposition of hCLDN18.2-zolbetuximab Fv (in dark blue) onto the structure of hCLDN18.2-osemitamab Fv (in light blue). (C) Left, the cryo-EM structure of the hCLDN18.2-zolbetuximab Fab complex. Zolbetuximab Fab is shown in surface representation, with the heavy chain colored in blue and the light chain in orange. The hCLDN18.2 structure is displayed in green cartoon. The binding interface between zolbetuximab Fab and hCLDN18.2 is indicated by a black dashed circle. Right, close-up view of the binding interface. CDRs of the heavy and light chains of zolbetuximab Fab are shown as blue and orange loops, respectively. ECL1 of hCLDN18.2 is shown in green cartoon. (D-G) Key interactions at the binding interface between zolbetuximab Fab and hCLDN18.2. Participating residues are displayed as sticks (D-G) or spheres (G). Potential hydrogen bonds are indicated by black dashed lines.

**Figure S3.**
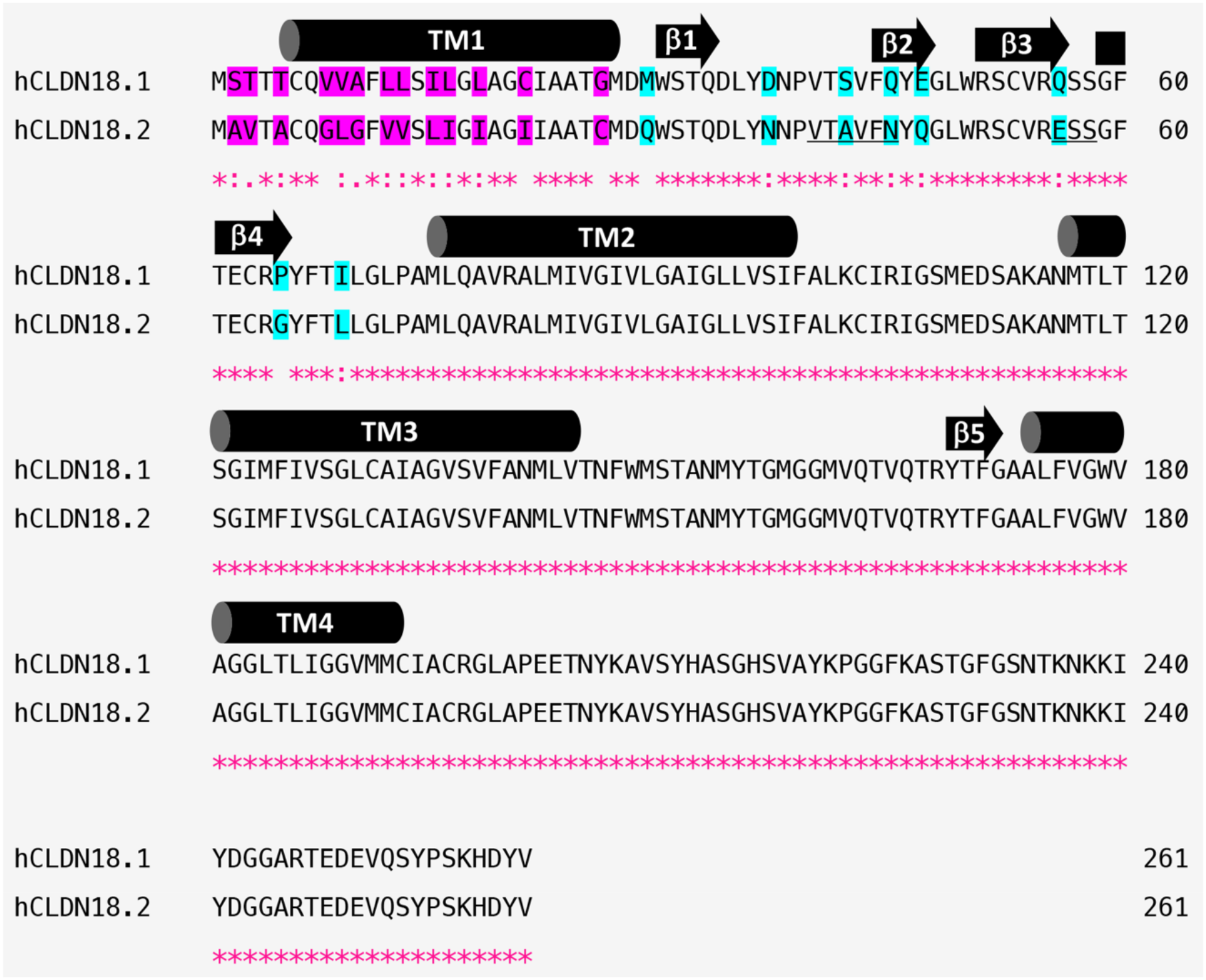
Sequence alignment of hCLDN18.1 and hCLDN18.2. Transmembrane helices are indicated by cylinders and labeled from TM1 to TM4, and β strands (β1-β5) are indicated by arrows. Residues that differ between the isoforms before or within TM1 are highlighted in purple, and different residues within ECL1 are highlighted in cyan. ICS1 (residues 40-45) and ICS2 (residues 56-58) of hCLDN18.2 are underlined. Asterisks (*) indicate identical residues. Colons (:) indicate strong similarities. Periods (.) indicate weak similarities.

**Figure S4.**
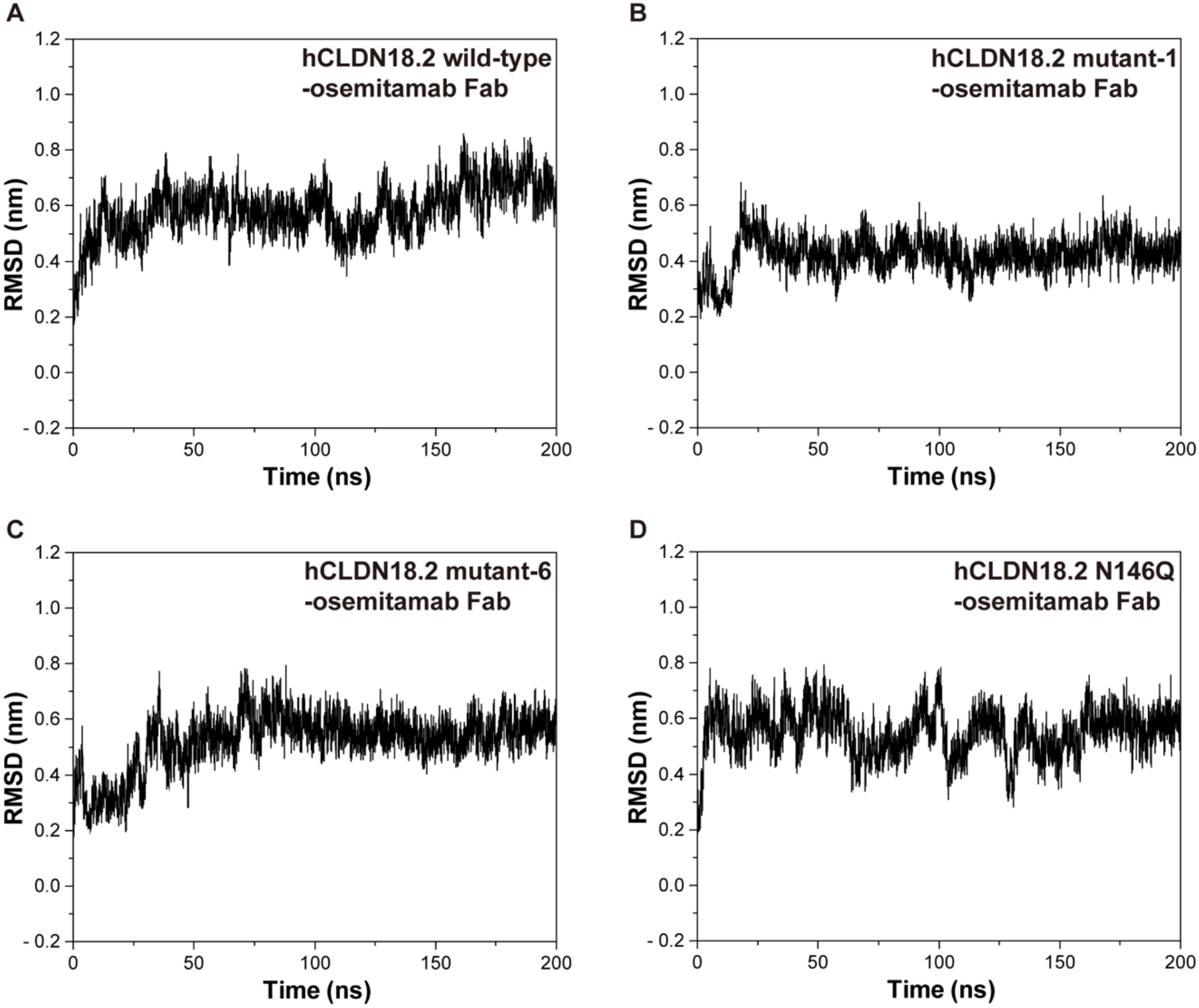
Backbone RMSD values of hCLDN18.2-osemitamab Fab complex (wild-type and mutants) during 200-ns MD simulations. (A) The complex of hCLDN18.2 wild-type and osemitamab Fab. (B) The complex of hCLDN18.2 mutant-1 and osemitamab Fab. (C) The complex of hCLDN18.2 mutant-6 and osemitamab Fab. (D) The complex of hCLDN18.2 N146Q and osemitamab Fab.

**Figure S5.**
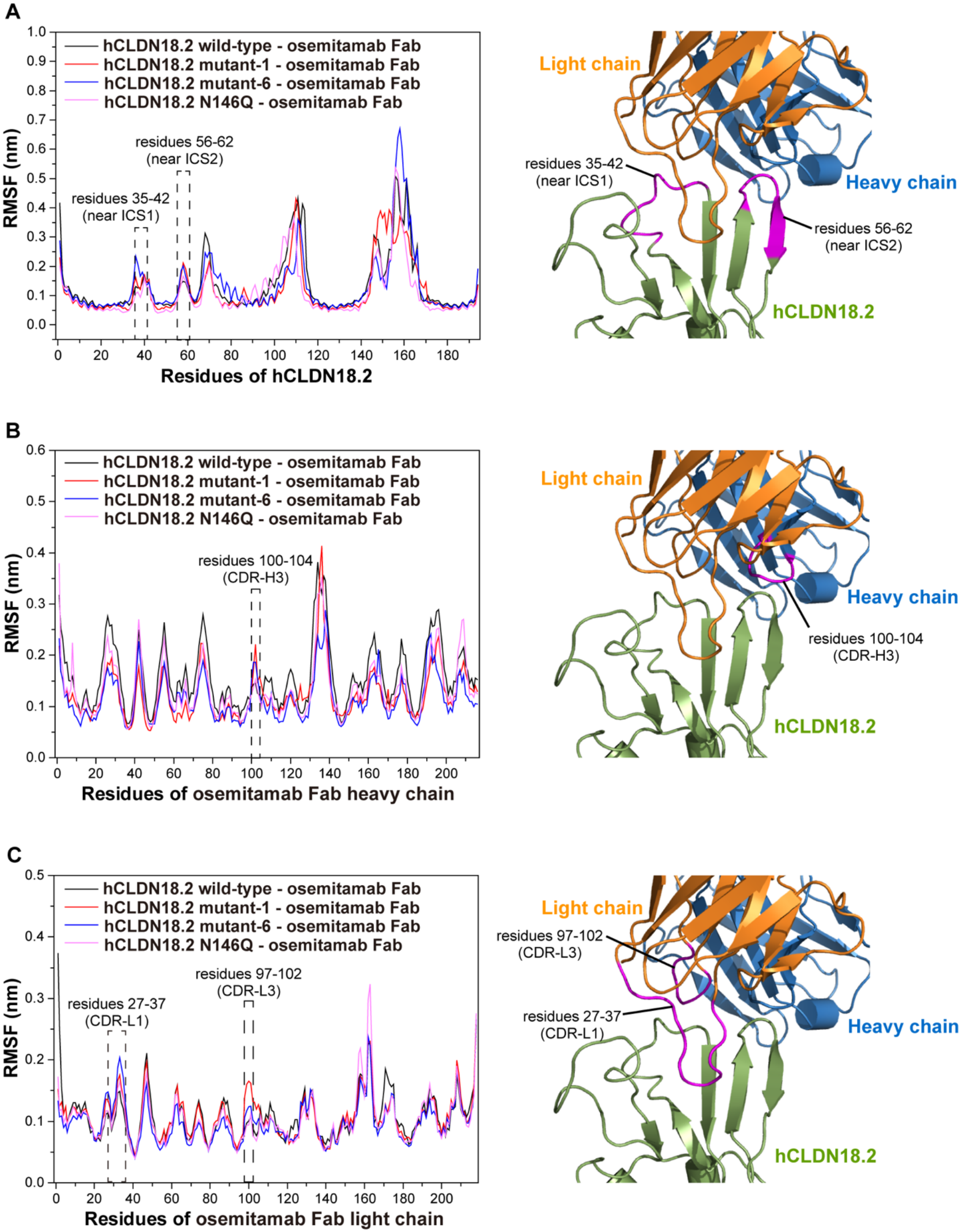
RMSF values of C_α_ atoms in hCLDN18.2-osemitamab Fab complex (wild-type and mutants). (A) RMSF analysis of hCLDN18.2 residues. (B) RMSF analysis of osemitamab Fab heavy chain residues. (C) RMSF analysis of osemitamab Fab light chain residues. Regions of interest are indicated by dashed rectangles in the left plots, and are displayed in magenta in the right cartoon.

**Figure S6.**
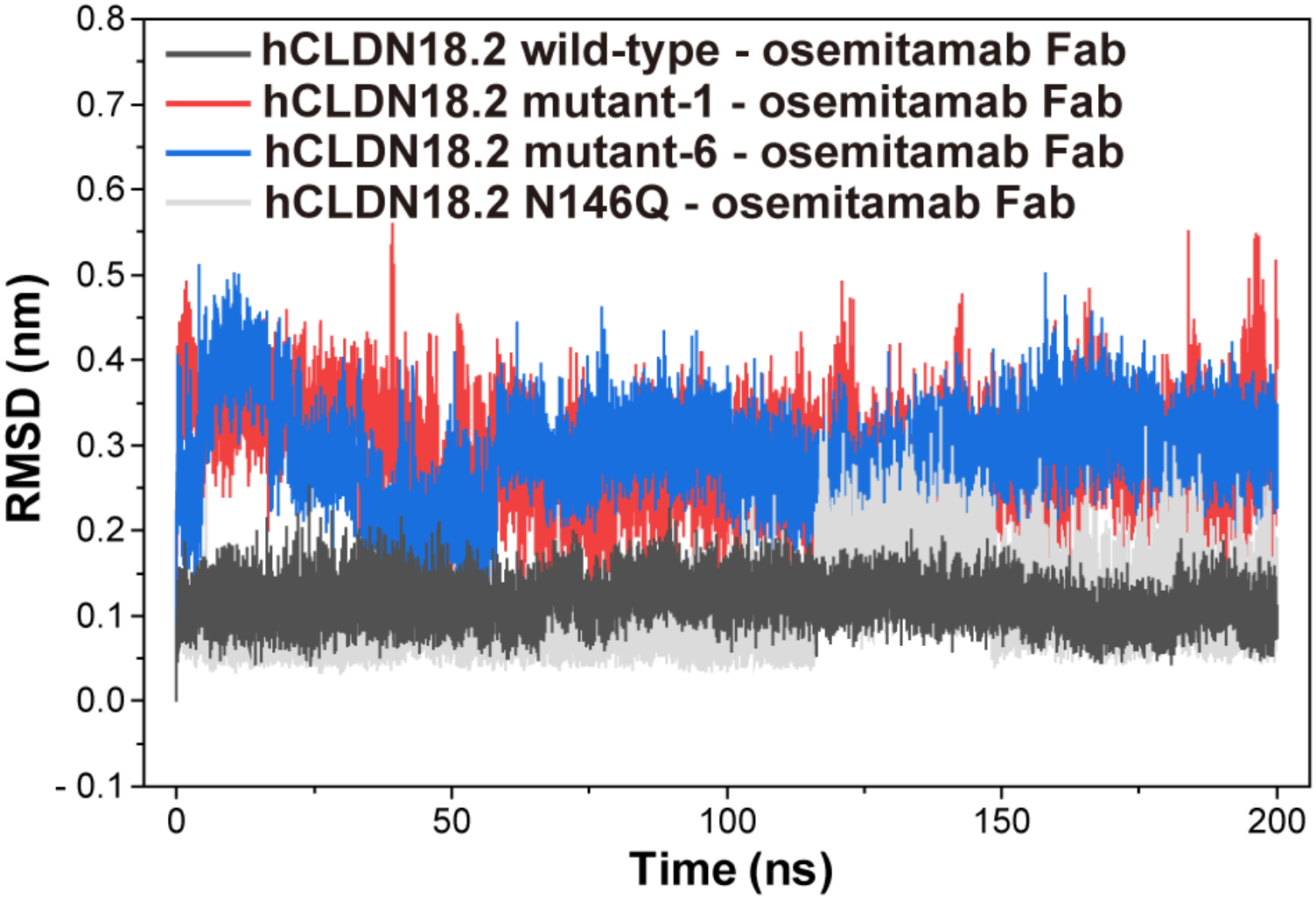
RMSD values of residues at non-bonded interacting sites between hCLDN18.2 wild-type and osemitamab Fab (listed in Table S3) during 200-ns MD simulations.

**Figure S7.**
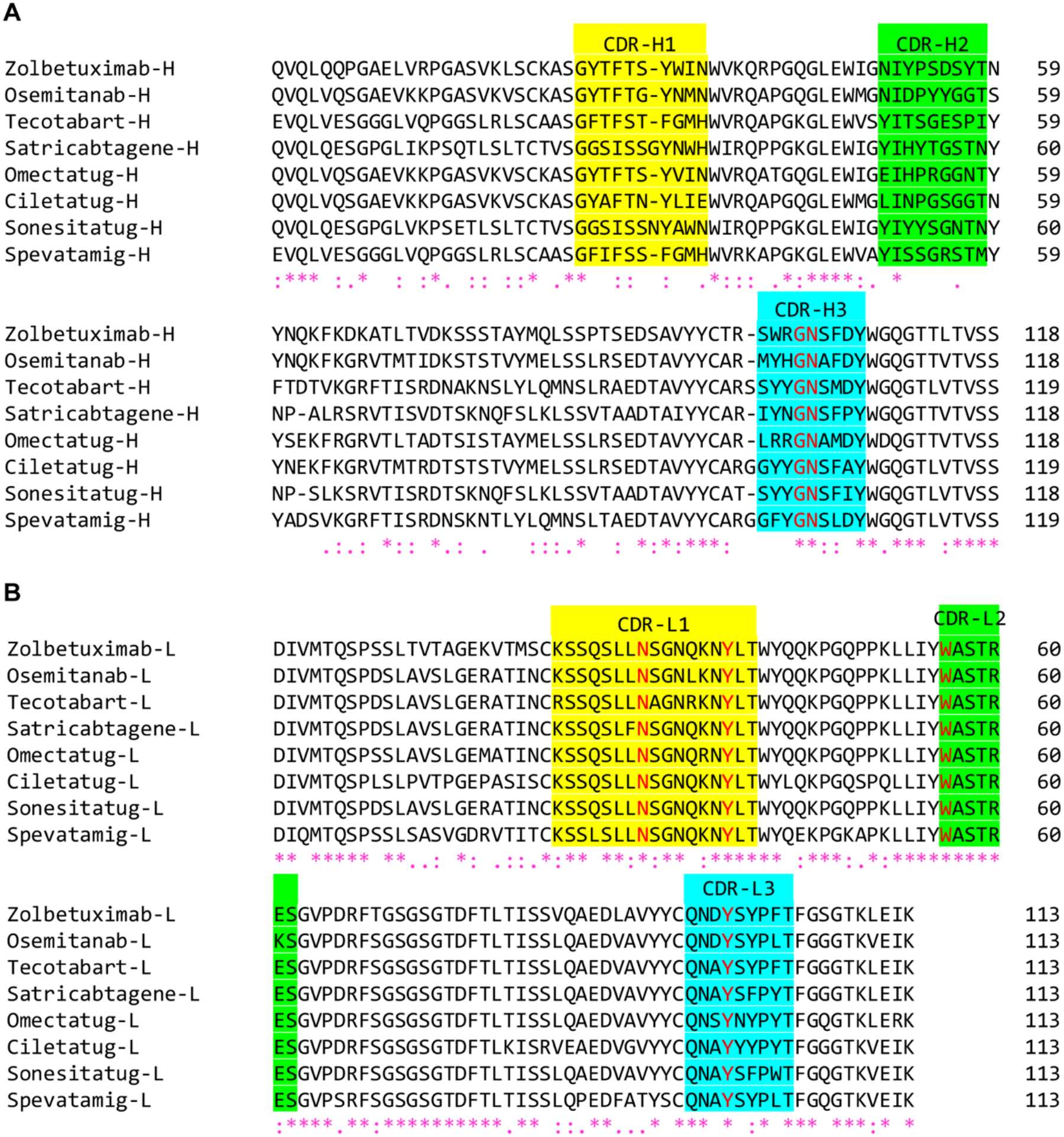
Sequence alignment of hCLDN18.2-targeting antibodies. Sequence alignment of heavy chain Fv regions (A) and light chain Fv regions (B) of eight hCLDN18.2-targeting antibodies. CDR-1, CDR-2, and CDR-3 are highlighted in yellow, green, and cyan, respectively. The CDR residues interacting with hCLDN18.2 residues Ala42, Asn45 and Glu56 in the hCLDN18.2-zolbetuximab/osemitamab Fab structures are shown in red. Asterisks (*) indicate identical residues. Colons (:) indicate strong similarities. Periods (.) indicate weak similarities.

**Figure S8.**
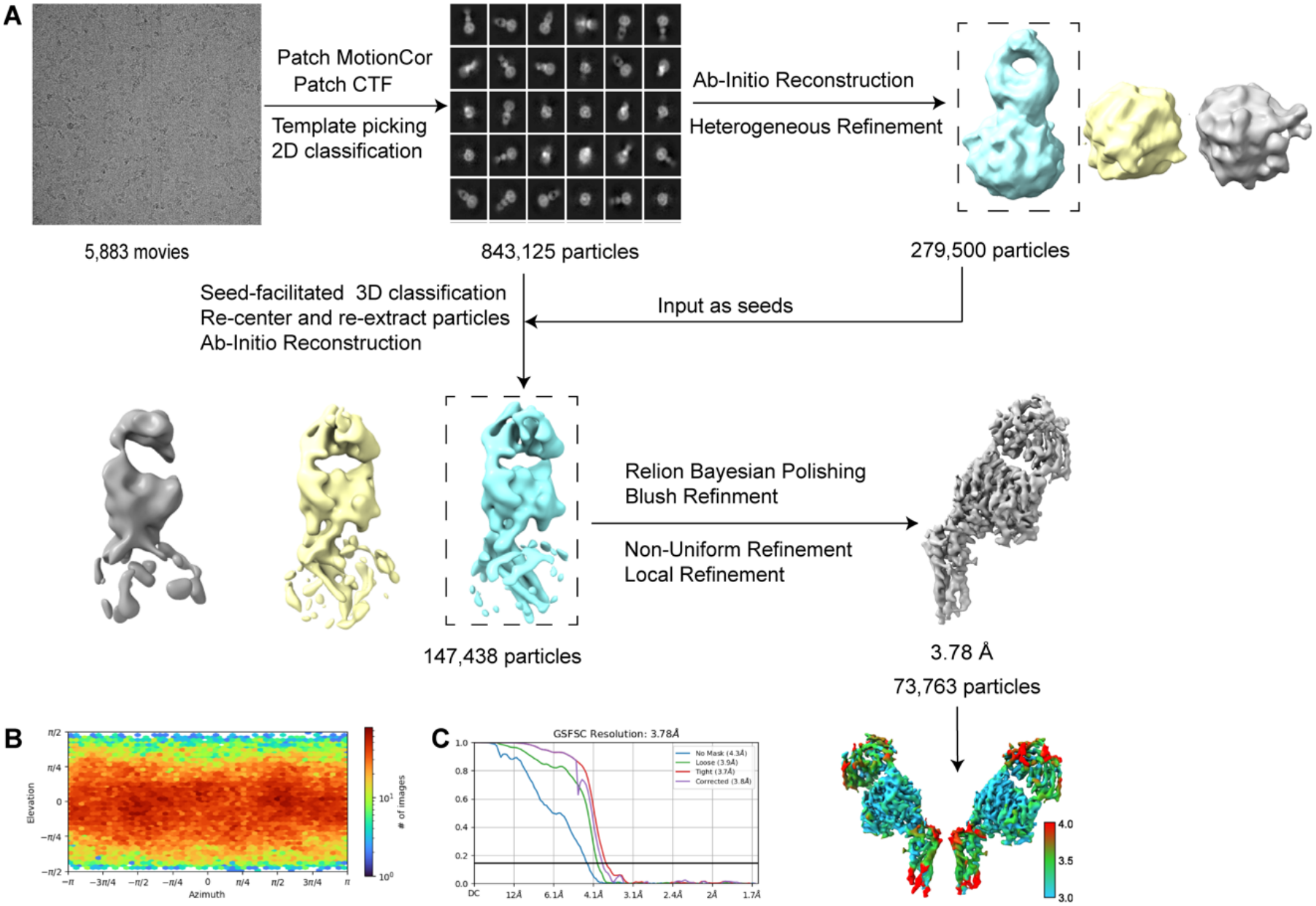
Cryo-EM data processing of monomeric hCLDN18.2-osemitamab Fab complex. (A) The workflow of cryo-EM data processing of the monomeric hCLDN18.2-osemitamab Fab complex. (B) Viewing direction distribution plot corresponding to C1-symmetry-expanded nonuniform refinement. (C) The overall resolution determined by the “gold standard” FSC curve using the FSC = 0.143 criterion.

**Figure S9.**
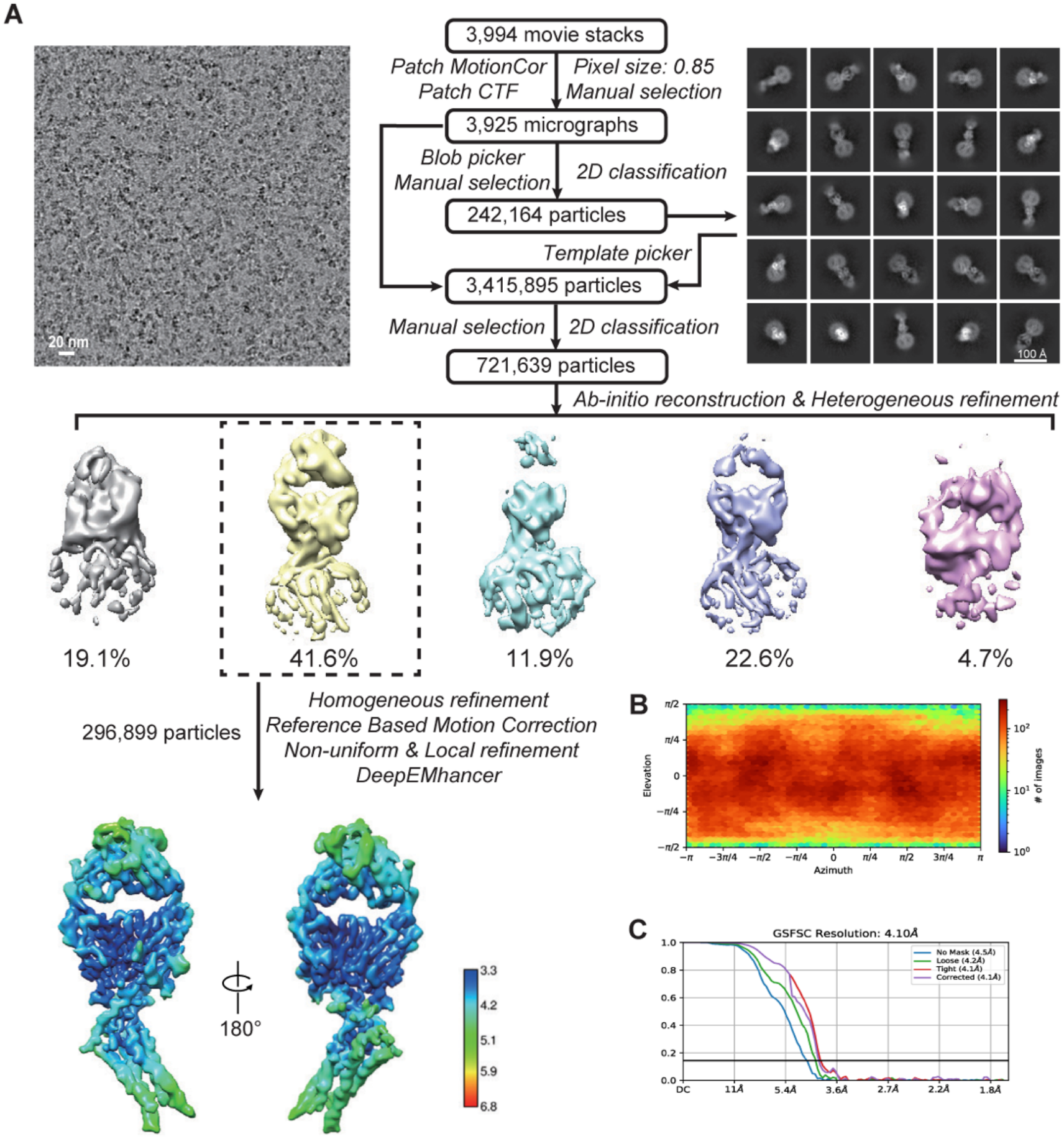
Cryo-EM data processing of monomeric hCLDN18.2-zolbetuximab Fab complex. (A) The workflow of cryo-EM data processing of the monomeric hCLDN18.2-zolbetuximab Fab complex. (B) Viewing direction distribution plot corresponding to C1-symmetry-expanded nonuniform refinement. (C) The overall resolution determined by the “gold standard” FSC curve using the FSC = 0.143 criterion.

**Figure S10.**
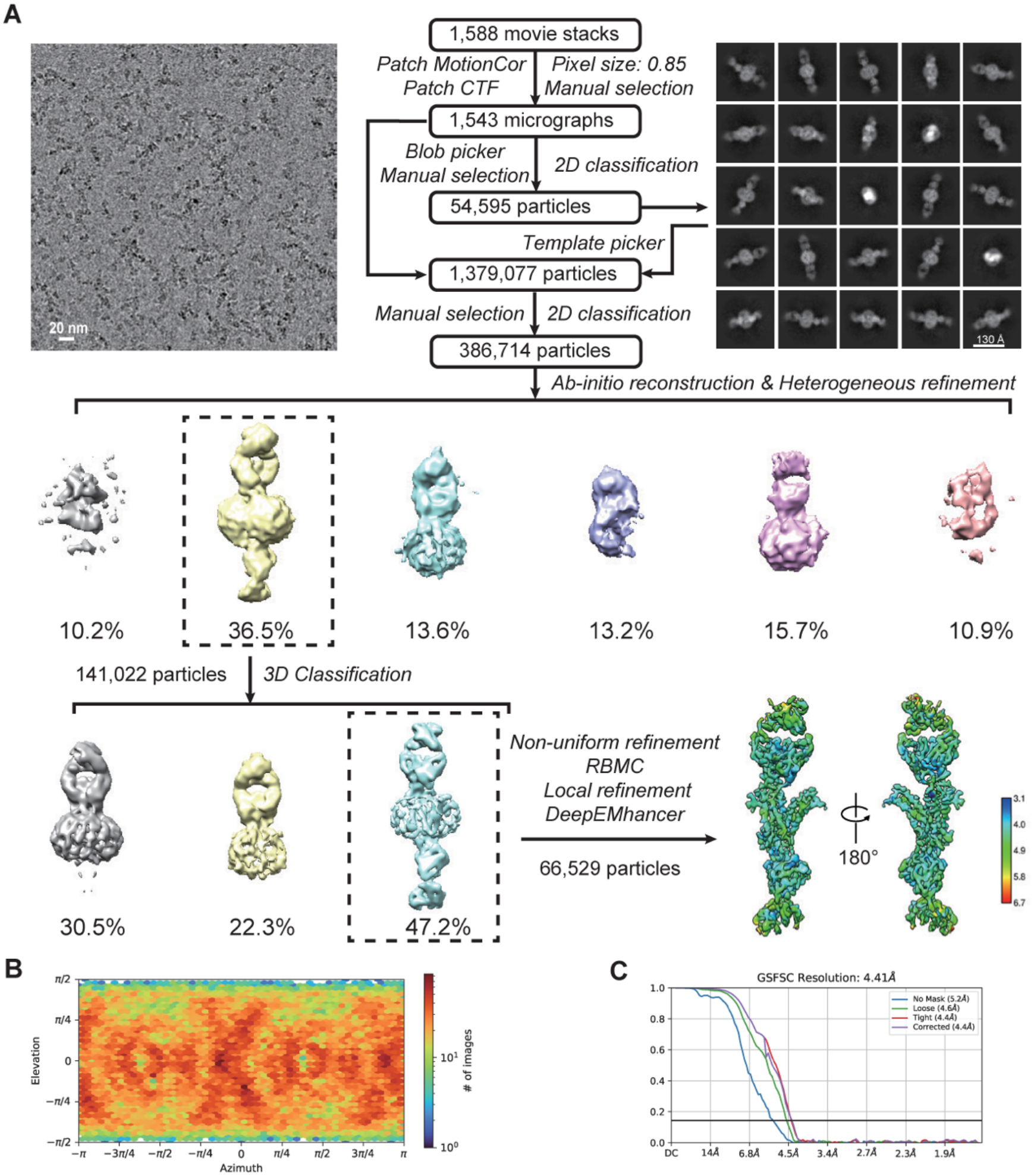
Cryo-EM data processing of dimeric hCLDN18.2-osemitamab Fab complex. (A) The workflow of cryo-EM data processing of the dimeric hCLDN18.2-osemitamab Fab complex. RBMC, reference-based motion correction. (B) Viewing direction distribution plot corresponding to C2-symmetry-expanded nonuniform refinement. (C) The overall resolution determined by the “gold standard” FSC curve using the FSC = 0.143 criterion.

**Table S1.** Cryo-EM data collection, refinement and validation statistics.

|  | **Monomeric hCLDN18.2-zolbetuximab Fab**<br>(EMDB-64747)<br>(PDB: 9V32) | **Monomeric hCLDN18.2-osemitamab Fab**<br>(EMDB-64740)<br>(PDB: 9V2U) | **Dimeric hCLDN18.2-osemitamab Fab**<br>(EMDB-64746)<br>(PDB: 9V31) |
|---|---|---|---|
| **Data collection and processing** |  |  |  |
| Movies | 3,994 | 5,883 | 1,588 |
| Magnification | 165,000 | 165,000 | 165,000 |
| Voltage (kV) | 300 | 300 | 300 |
| Electron exposure (e<sup>–</sup>/Å<sup>2</sup>) | 57.5 | 56.4 | 57.5 |
| Defocus range (μm) | -1.0 to -1.8 | -0.1 to -3 | -1.0 to -1.8 |
| Pixel size (Å) | 0.85 | 0.85 | 0.85 |
| Symmetry imposed | C1 | C1 | C2 |
| Initial particle images (no.) | 3,415,895 | 843,125 | 1,379,077 |
| Final particle images (no.) | 296,899 | 73,763 | 66,529 |
| Map resolution (Å) | 4.1 | 3.8 | 4.4 |
| FSC threshold | 0.143 | 0.143 | 0.143 |
| Map sharpening *B* factor (Å<sup>2</sup>) | -206.6 | -196.6 | -197.8 |
| **Refinement** |  |  |  |
| Model composition |  |  |  |
| Non-hydrogen atoms | 4,507 | 4,380 | 8,839 |
| Protein residues | 593 | 577 | 1,165 |
| Nucleotides | - | - | - |
| Ligands | - | - | - |
| *B* factors (Å<sup>2</sup>, mean) |  |  |  |
| Protein | 82.91 | 67.70 | 71.12 |
| Nucleotide | - | - | - |
| Ligand | - | - | - |
| R.m.s. deviations |  |  |  |
| Bond lengths (Å) | 0.003 | 0.003 | 0.003 |
| Bond angles (°) | 0.637 | 0.675 | 0.658 |
| Validation |  |  |  |
| MolProbity score | 1.75 | 1.64 | 1.70 |
| Clashscore | 9.60 | 9.77 | 10.81 |
| Poor rotamers (%) | 0.20 | 0.00 | 0.00 |
| Ramachandran plot |  |  |  |
| Favored (%) | 96.37 | 97.33 | 97.18 |
| Allowed (%) | 3.63 | 2.67 | 2.74 |
| Disallowed (%) | 0.00 | 0.00 | 0.09 |

**Table S2.**
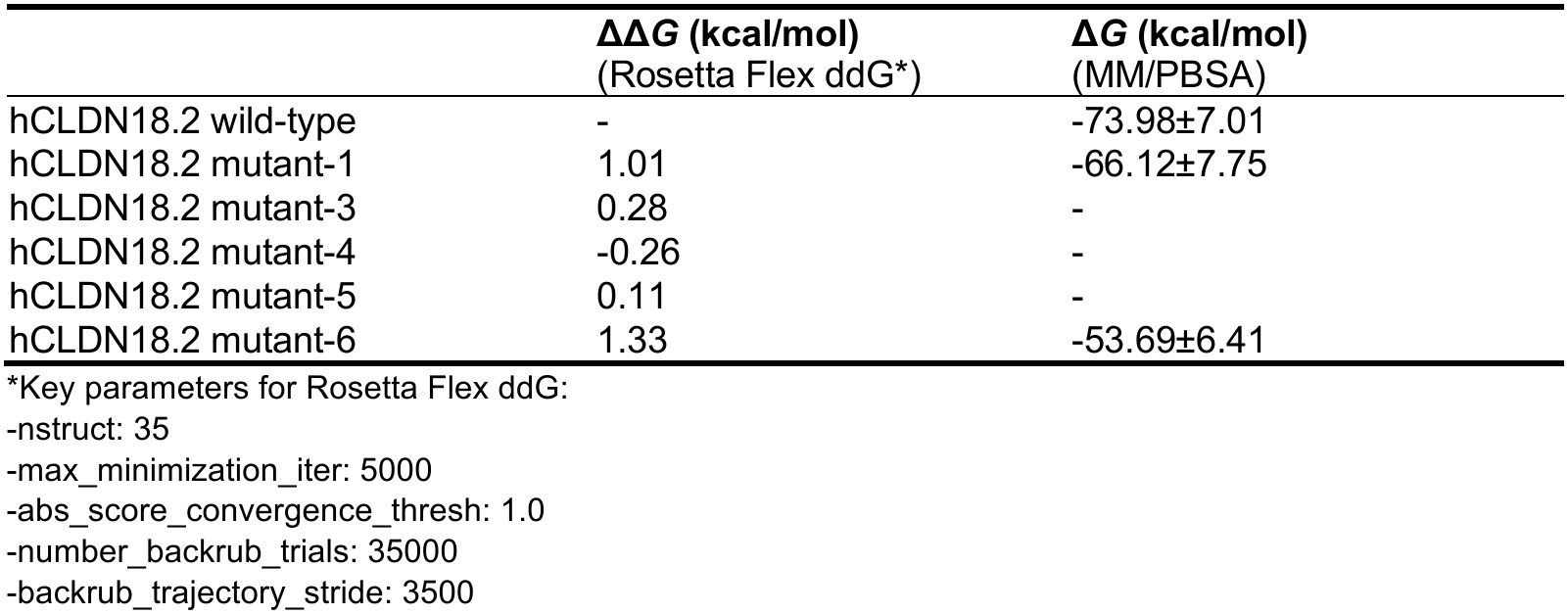
ΔΔ*G* and Δ*G* of osemitamab Fab binding to hCLDN18.2 variants.

**Table S3.** Non-bonded interactions between osemitamab Fab and hCLDN18.2 during MD.

| **hCLDN18.2** | **Osemitamab Fab** | **Distance (Å)** | **Non-bonded category** |
|---|---|---|---|
| Chain A*:Glu56:OE1 | Chain B:Gly102:H | 1.9 | Hydrogen bond |
| Chain A:Ser57:HG | Chain B:Asn103:OD1 | 2.0 | Hydrogen bond |
| Chain A:Ser58:H | Chain B:Asn103:OD1 | 2.7 | Hydrogen bond |
| Chain A:Ala42:O | Chain B:Asn103:HD21 | 2.8 | Hydrogen bond |
| Chain A:Glu62:OE1 | Chain C:Lys36:HZ2 | 1.8 | Hydrogen bond; Electrostatic |
| Chain A:Glu56:OE2 | Chain C:Trp56:HE1 | 1.9 | Hydrogen bond |
| Chain A:Glu62:OE2 | Chain C:Asn34:HD22 | 2.0 | Hydrogen bond |
| Chain A:Gln163:HE21 | Chain C:Tyr98:OH | 2.0 | Hydrogen bond |
| Chain A:Thr164:OG1 | Chain C:Ser32:HB2 | 2.5 | Hydrogen bond |
| Chain A:Glu56:OE2 | Chain C:Tyr38:HH | 2.8 | Hydrogen bond |
| Chain A:Ala42:H | Chain C:Tyr98:O | 3.0 | Hydrogen bond |
| Chain A:Ala42 | Chain C:Tyr100 | 4.6 | Hydrophobic |
| Chain A:Val43 | Chain C:Tyr38 | 4.8 | Hydrophobic |

\*Chain A: hCLDN18.2; Chain B: osemitamab Fab heavy chain; Chain C: osemitamab Fab light chain.

